# CRISPR-Mediated Targeting of BRAF Oncogenes in Pediatric Low-Grade Glioma

**DOI:** 10.64898/2026.08.12.744431

**Authors:** Christy A. George, Margaret E. Brown, Priya Rana, Deirdre A. Killebrew, Ross C. Wilson

## Abstract

**Summary:** A catch-all intronic guide RNA pair excises the KIAA1549—BRAF oncofusion across its major variants, with productive junction excision confirmed by gain-of-function PCR in patient-derived glioma cells. An allele-specific guide selectively disrupts BRAF V600E, in patient-derived pediatric low-grade glioma cells.

Pediatric low-grade glioma (pLGG) is the most common brain tumor of childhood, accounting for 30—50% of all pediatric central nervous system malignancies^1^. The disease is almost universally driven by activating mutations in the BRAF serine/threonine kinase: a chromosomal tandem duplication generating the KIAA1549—BRAF oncofusion in approximately 70% of cases, or the BRAF V600E gain-of-function point mutation in approximately 15%^2^. Current targeted pharmacotherapies, including the RAF inhibitor tovorafenib, require continuous dosing, are not allele-specific, and carry risks of long-term toxicity in children. A one-time genomic intervention that permanently disables the oncogenic BRAF alteration while preserving wild-type BRAF signaling represents a compelling therapeutic alternative. In this study, we describe the design and experimental validation of allele-specific CRISPR guide RNAs targeting both the KIAA1549—BRAF oncofusion and the BRAF V600E point mutation. For the oncofusion, we developed a double-cut intronic excision strategy in which a guide RNA targeting KIAA1549 intron 14 is paired with a guide RNA targeting BRAF intron 11. Because the genomic breakpoints of all four major fusion variants (KB 16:9, 15:9, 16:11, and 15:11) fall within these introns, a single guide pair can address the full landscape of fusion heterogeneity in a single intervention. For BRAF V600E, we exploited a unique PAM sequence created by the pathogenic TBA transversion at codon 600, enabling allele-specific SpCas9 and AsCas12a guide designs that distinguish the mutant from the wild-type allele at single-nucleotide resolution. We screened guide RNA candidates by ribonucleoprotein (RNP) nucleofection in A375 human melanoma cells (BRAF V600E homozygous) and in patient-derived 3635 PXA glioma cells (BRAF V600E heterozygous). The top KIAA1549 intron 14 guide, K9_i14_A_Cas9, achieved 66% indel frequency in A375 cells. The top BRAF intron 11 guides, B_i11_A_Cas9 and B_i11_D_Cas9, achieved 84% and 85% indel frequency, respectively. For BRAF V600E, the best allele-specific SpCas9 guide achieved l57% editing in A375 cells and l74% editing in 3635 PXA patient-derived glioma cells. Dual-cut excision of the KIAA1549—BRAF junction was confirmed by a gain-of-function PCR assay designed to detect the excision junction amplicon (∼191 bp) produced by NHEJ-mediated rejoining of the KIAA1549 intron 14 and BRAF intron 11 cut ends.

## INTRODUCTION

Pediatric low-grade glioma (pLGG) encompasses a heterogeneous group of World Health Organization (WHO) grade I and II central nervous system tumors arising in children and adolescents, accounting for 30% of all pediatric brain tumors with an annual incidence of approximately 2—3 per 100,000 children^1,3^. The clinical spectrum spans from circumscribed, surgically resectable tumors — most commonly pilocytic astrocytoma (PA, WHO grade I), which carries an excellent 10-year survival exceeding 90% following gross total resection — to diffuse, infiltrative tumors involving eloquent brain structures such as the optic pathway, hypothalamus, brainstem, and spinal cord, where complete resection is impossible and long-term morbidity is high^1,4^.

The management of unresectable or progressive pLGG has evolved significantly over the past decade but remains unsatisfactory. Conventional chemotherapy (carboplatin/vincristine) produces transient responses in the majority of patients but is associated with significant toxicity and high rates of relapse after treatment cessation^5^. Radiation therapy, while effective for local tumor control, causes devastating neurocognitive sequelae in young children owing to the broad fields required and the vulnerability of the developing brain, and is generally reserved for older adolescents or cases refractory to all other modalities^6^. The advent of molecularly targeted therapies — specifically MEK inhibitors (selumetinib, trametinib) and the RAF inhibitor tovorafenib (approved by the FDA in 2024 for relapsed or refractory BRAF-altered pLGG) — has transformed the treatment landscape and produces durable disease control in a substantial fraction of patients^7—9^. However, these agents require continuous daily or weekly dosing for an indefinite period, their long-term toxicity profiles in children remain under investigation, and, importantly, they do not address the underlying oncogene at the genomic level. A single treatment that permanently disables the driving mutation could, in principle, obviate the need for long-term pharmacotherapy and the associated risks.

The defining molecular feature of pLGG at the cellular level is activation of the mitogen-activated protein kinase (MAPK) signaling cascade. The MAPK pathway operates through the sequential kinase relay RAS → RAF → MEK → ERK, which regulates cell proliferation, differentiation, survival, and senescence in response to growth factor stimulation. In normal cells, MAPK signaling is tightly regulated through feedback mechanisms, receptor-level desensitization, and the intrinsic GTPase activity of RAS. In pLGG, activating alterations in BRAF, the predominant RAF isoform active in neural precursor cells, constitutively activate this pathway independent ofupstream growth factor signals, driving tumor cell proliferation^10^.

BRAF encodes a serine/threonine kinase that functions in the MAPK cascade as a signal transducer downstream of RAS. In the basal state, BRAF is autoinhibited through an intramolecular interaction between its N-terminal regulatory domain (containing the RAS-binding domain, or RBD) and its C-terminal kinase domain. RAS-GTP binding to the RBD releases this autoinhibition, enabling BRAF dimerization and kinase activation. Both major pLGG-associated alterations disrupt this regulation: the KIAA1549—BRAF fusion removes the N-terminal regulatory domain entirely, while the BRAF V600E substitution mimics the activated conformation and confers constitutive kinase activity independent of RAS input^11^.

The BRAF V600E mutation arises from a single nucleotide transversion at codon 600 of the BRAF coding sequence: a thymine-to-adenine change at cDNA position 1799 (c.1799T>A) that substitutes glutamic acid for valine at residue 600 (p.Val600Glu)^10^. This substitution inserts a negatively charged residue into the activation loop of the kinase domain, mimicking the phosphorylated state that normally follows RAS stimulation and locking BRAF in a constitutively active conformation. The resulting kinase exhibits approximately 500-fold elevated activity compared to wild-type BRAF and is capable ofactivating MEK as a monomer, bypassing the requirement for RAS-GTP and BRAF dimerization^10^. BRAF V600E is present in approximately 15% of pLGG cases overall and is enriched in specific histological subtypes including pleomorphic xanthoastrocytoma (PXA), diffuse astrocytoma, and ganglioglioma^12^.

The single-nucleotide nature of the V600E mutation creates a specific opportunity for allele-specific genome editing that exploits the PAM requirement of CRISPR-Cas systems. SpCas9 requires a 5’-NGG-3’ protospacer adjacent motif (PAM) immediately 3’ of the target site for binding and cleavage^13^. The c.1799T>A transversion in V600E creates a novel NGG PAM sequence on the non-template strand that is absent in the wild-type BRAF sequence, where a TGG motif is replaced by AGG in the mutant context. A guide RNA whose spacer places the mutant adenine residue at position 3 from the PAM (the position most sensitive to mismatches) can distinguish the V600E from the wild-type allele with high specificity: the NGG PAM is present only on the mutant allele, and wild-type BRAF cells will not be cut because the PAM is absent. An analogous allele-specific opportunity exists for AsCas12a, which requires a 5’-TTTV-3’ PAM^14^: the V600E transversion creates a novel TTTV PAM on the non-template strand that is not present in wild-type BRAF.

The KIAA1549—BRAF (KB) oncofusion is generated by a tandem duplication of approximately 2 megabases on chromosome 7q34 (hg38: approximately chr7:138,400,000—140,900,000) that juxtaposes the KIAA1549 gene (encoding a protein of unknown function with predicted transmembrane domains) upstream of BRAF in head-to-tail orientation^15^. The resulting fusion protein retains the intact BRAF kinase domain (C-terminus) fused to the N-terminal and middle segments of KIAA1549, from which the N-terminal regulatory receptor binding domain of BRAF is absent. This architectural loss of autoregulation constitutively activates the kinase domain and drives downstream MEK—ERK signaling independent of RAS input.

Four major KIAA1549—BRAF fusion variants have been described, distinguished by the exon-level boundaries at which the two genes are joined: KB 16:9 (KIAA1549 exon 16 fused to BRAF exon 9), KB 15:9 (KIAA1549 exon 15 to BRAF exon 9), KB 15:11 (KIAA1549 exon 15 to BRAF exon 11), and KB 16:11 (KIAA1549 exon 16 to BRAF exon 11). The KB 16:9 variant is most prevalent, accounting for approximately 78% of all KB fusions; together, the four variants represent the vast majority of KB fusion-positive pLGG^2^. Despite their different exon boundaries, all four fusion variants share a critical structural property: the chromosomal breakpoints giving rise to the tandem duplication fall not within exonic sequences but within intronic regions of both genes. Specifically, the genomic breakpoints of all four major variants fall within KIAA1549 intron 14 or intron 15, and within BRAF intron 9 or intron 11 ^16,17^.

This convergence of intronic breakpoints across all four fusion variants underlies the therapeutic strategy described here. Rather than targeting the unique fusion junction sequence, which differs at the nucleotide level for each variant and would require patient-specific guide design, we targeted intronic sequences that are common to all variants. Guide RNAs that cut within KIAA1549 intron 14 (present in all KB fusions regardless of whether the exon 15 or exon 16 boundary is used) and within BRAF intron 11 (present in both KB 16:11 and in very close proximity to intron 9 breakpoints of KB 16:9 and 15:9) form a pair that, when both cuts are executed, excises the duplicated chromosomal segment and enables NHEJ-mediated rejoining of the flanking intact chromosomal ends. The resulting repaired chromosome lacks the duplicated segment and encodes no fusion protein, thereby enabling a permanent, one-cut genomic correction applicable to all four major KB fusion variants simultaneously.

The intronic breakpoint architecture and allele-specific PAM opportunities make pLGG an unusually tractable target for CRISPR-based therapy. Unlike many solid tumors, in which multiple redundant oncogenic drivers and tumor suppressor losses complicate single-gene targeting, pLGG is defined by a single dominant activating alteration in the vast majority of cases. Disruption of this dominant alteration is expected to be sufficient for therapeutic effect based on established oncogene addiction biology.

Current targeted therapies (tovorafenib, selumetinib) inhibit BRAF or MEK kinase activity at the protein level and are effective, but they do not distinguish between the oncogenic and wild-type BRAF proteins. This non-specificity creates two clinical problems: first, RAF inhibitor monotherapy can paradoxically activate the MAPK pathway in cells expressing wild-type BRAF by promoting RAS-independent BRAF dimerization^18,19^, contributing to adverse effects including skin toxicity and potential secondary malignancies; and second, continuous pharmacological pressure selects for resistance mechanisms that re-activate MEK—ERK signaling downstream of the drug target. CRISPR editing of the oncogenic allele specifically addresses both of these concerns. By targeting intronic sequences present only in the duplicated chromosomal segment (for the KB fusion) or a PAM that exists only on the mutant allele (for V600E), the edit is restricted to the oncogenic allele, leaving wild-type BRAF intact. A permanent genomic disruption also fundamentally forecloses certain resistance mechanisms, such as re-expression of the oncogene through transcriptional upregulation, which can circumvent kinase inhibition.

A two-cut excision of approximately 30 kilobases — the largest possible distance between KIAA1549 intron 14 and BRAF intron 11 in the duplicated chromosomal segment — is precisely the type of coordinated dual-cut editing for which synchronized guide RNA expression is most important.

## RESULTS

### Development of a genome editing strategy for KIAA1549—BRAF

We designed the KIAA1549—BRAF editing strategy around two key principles. First, to address all four major KB fusion variants with a single guide pair, target sites must lie within intronic sequences present in all variants — not within the fusion junction itself, which differs in primary sequence among variants. Second, both cuts must fall within the duplicated chromosomal region of the fusion gene such that NHEJ-mediated repair results in a programmed excision. Because the WT copies of BRAF and KIAA1549 are separated by nearly 2 megabases, we anticipate that intronic cuts in the WT genes will not contribute to programmed excisions. Thoughtfully designed gRNAs targeting each intron are expected to result in indels that do not impact splicing and thus protect WT gene function.

The tandem duplication at 7q34 that generates the KB fusion brings the 3’ portion of KIAA1549 into proximity with the 5’ portion of BRAF. In the WT chromosome, KIAA1549 and BRAF are separated by approximately 1.9 megabases on 7q34. After the duplication, a head-to-tail copy of this segment is inserted, placing a copy of the BRAF 5’ end (including introns 9 and 11) adjacent to a copy of the KIAA1549 3’ end (including introns 14 and 15). The breakpoints that define each fusion variant fall within these intronic regions. A pair of guide RNAs cutting within KIAA1549 intron 14 (present upstream of the breakpoint in KB 16:9 and 15:9 variants) and within BRAF intron 11 (present downstream of the breakpoint in KB 16:11 and 15:11 and in very close genomic proximity to intron 9 breakpoints in KB 16:9 and 15:9) will, when both cuts are made, excise the pathogenic duplicated segment.

### Identification of intronic target sites

Candidate guide RNA spacers within KIAA1549 intron 14 and BRAF intron 11 were identified by submitting each intron sequence to CRISPOR^20^ against the human reference genome (hg38), using SpCas9 (NGG PAM) as the primary search enzyme and AsCas12a (TTTV PAM) for secondary V600E allele-specific design. For each candidate, we computed the Doench 2016 on-target activity score and the MIT specificity score, which reflects the weighted off-target profile across the genome with up to four mismatches.

KIAA1549 intron 14 (hg38: chr7:138,869,537-138,868,129; 1, 408 nucleotides; reverse strand) was selected as the left-guide target region because it is the shared intronic sequence common to all four major KB fusion variants on the KIAA1549 side of the breakpoint. A panel of candidate guides within this intron was synthesized and designated with the prefix K9_i14 (KIAA1549 intron 14) followed by a letter identifier. BRAF intron 11 (hg38: chr7:140,783,020-140,781, 694; 1, 326 nucleotides) was selected as the right-guide target region, as it is present downstream of the BRAF intron 11 breakpoint and lies within the duplication segment relevant to all four variants. Guides within this region were designated B_i11.

For BRAF V600E, the allele-specific strategy exploits the c.1799TBA transversion. On the coding strand, the wild-type sequence at codon 600 is 5’-GTG-3’ (valine); in the V600E mutant, this is 5’-GAG-3’ (glutamic acid). On the non-template strand, the wild-type sequence at this position is 5’-CAC-3’; the V600E sequence is 5’-CTC-3’. Inspection of the flanking sequences revealed that the TBA change at position 1799 creates an AGG PAM on the non-template strand in the V600E context (where wild-type has a TCG sequence lacking a suitable PAM), enabling allele-specific targeting by SpCas9. The same transversion also creates a TTTTC PAM variant compatible with AsCas12a on the other strand. Guide RNA spacers were designed such that the single nucleotide distinguishing V600E from wild-type falls within the seed region ofthe spacer (positions 14-20 from the PAM-distal end), maximizing allele discrimination.

To ensure that editing at the intronic target sites would not disrupt known regulatory elements or coding sequences at off-target loci, all candidate guides were screened by CRISPOR for predicted off-target sites with up to four mismatches in the human genome. Candidates with predicted high-confidence off-targets (two or fewer mismatches) in exonic sequences of any protein-coding gene were excluded from further evaluation. The remaining candidates, meeting criteria for on-target score, off-target specificity, and intronic positioning relative to the fusion breakpoint architecture, were advanced to experimental validation.

### Computational assessment of on-target and off-target activity

*In silico* evaluation of all candidate guides produced a ranked list of spacers based on combined on-target activity prediction and off-target specificity. For KIAA1549 intron 14 candidates, on-target scores ranged from 17-66% across the candidate panel, with the K9_i14_A spacer achieving the highest on-target score and a favorable off-target profile. For BRAF intron 11 candidates, B_i11 _A and B_i11 _D achieved high predicted on-target activity with no predicted off-targets with fewer than three mismatches in exonic sequences of the hg38 genome.

For the BRAF V600E allele-specific guides, *in silico* analysis confirmed that the mutant-specific NGG PAM recognized by the SpCas9-targeted spacer is absent in the wild-type BRAF sequence at codon 600. The single nucleotide difference between wild-type (T at position 1799) and mutant (A at position 1799) falls within the seed region of the designed spacer — the 7 nucleotides immediately 5’ of the PAM — where mismatches are most likely to ablate Cas9 activity^21^. This positioning provides the primary basis for allele specificity: even if occasional off-target binding occurs at the wild-type locus, the mismatch at a seed-region position is expected to substantially impair cleavage efficiency at the wild-type allele.

A full table of guide sequences, CRISPOR scores, predicted off-target loci, and allele-specificity assessment is provided in Table 1. The three guides selected for advancement to experimental validation were K9_i14_A (KIAA1549 intron 14), B_i11 _A and B_i11 _D (BRAF intron 11), and the lead V600E allele-specific SpCas9 guide.

**Table 1.**
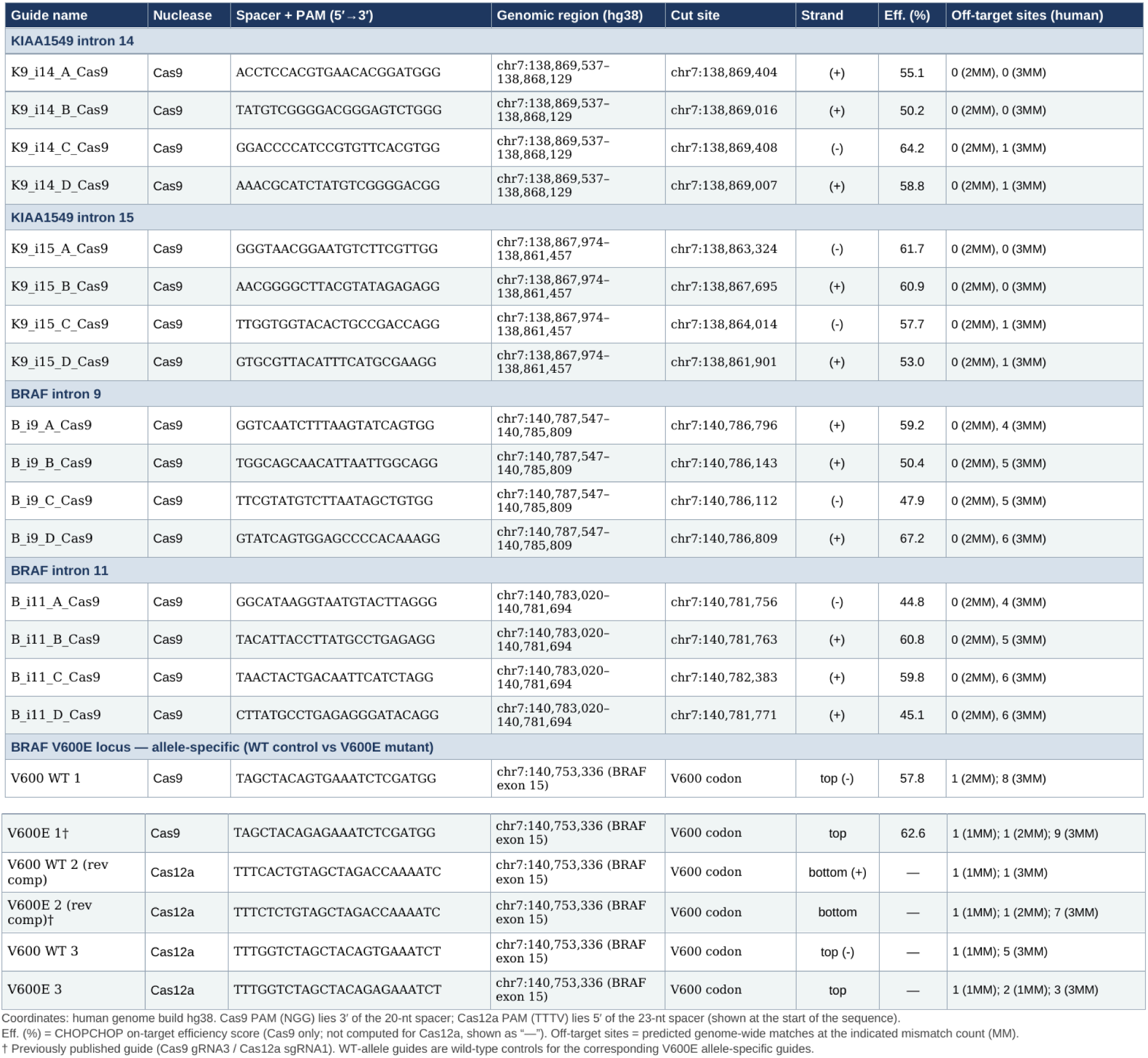
Candidate BRAF V600E and KIAA1549—BRAF intronic sgRNAs.

### Validation of sgRNA activity

Candidate guide RNAs were screened for on-target editing efficiency by ribonucleoprotein (RNP) nucleofection in two cell lines: A375 human melanoma cells, which carry a homozygous BRAF V600E mutation and provide a readily transfectable reference line for both the V600E allele-specific guides and the intronic excision guides; and 3635 PXA patient-derived pleomorphic xanthoastrocytoma cells, which carry a heterozygous BRAF V600E mutation in a CNS tumor cell background more representative of the target disease.

RNP complexes were assembled from IDT Alt-R sgRBA annealed, complexed with a 0C-triNLS SpCas9 protein manufactured by Aldevron, at a 1.2:1 sgRNA:Cas9 molar ratio. RNPs were delivered to cells by nucleofection (Lonza SF 4D-Nucleofector), and cells were harvested 48-72 hours later for genomic DNA extraction and ICE analysis. Indel frequency at each target site was quantified from Sanger sequencing traces of PCR amplicons flanking each cut site, analyzed using Synthego ICE v3.

Screening of KIAA1549 intron 14 candidates in A375 cells identified K9_i14_A_Cas9 as the most active guide, achieving 66% indel frequency by ICE (Fig. 3). Screening of BRAF intron 11 candidates identified B_i11 _A_Cas9 (84% indel frequency) and B_i11 _D_Cas9 (85% indel frequency) as the top performers in A375 cells, with B_i11 _D_Cas9 selected as the primary lead based on its slightly higher activity and favorable CRISPOR specificity profile (Fig. 3). The high editing efficiencies achieved at BRAF intron 11 — consistently above 80% — likely reflect the favorable chromatin environment and sequence context of this intronic region, which has proven accessible to Cas9 in multiple cell types.

**Figure 1.**
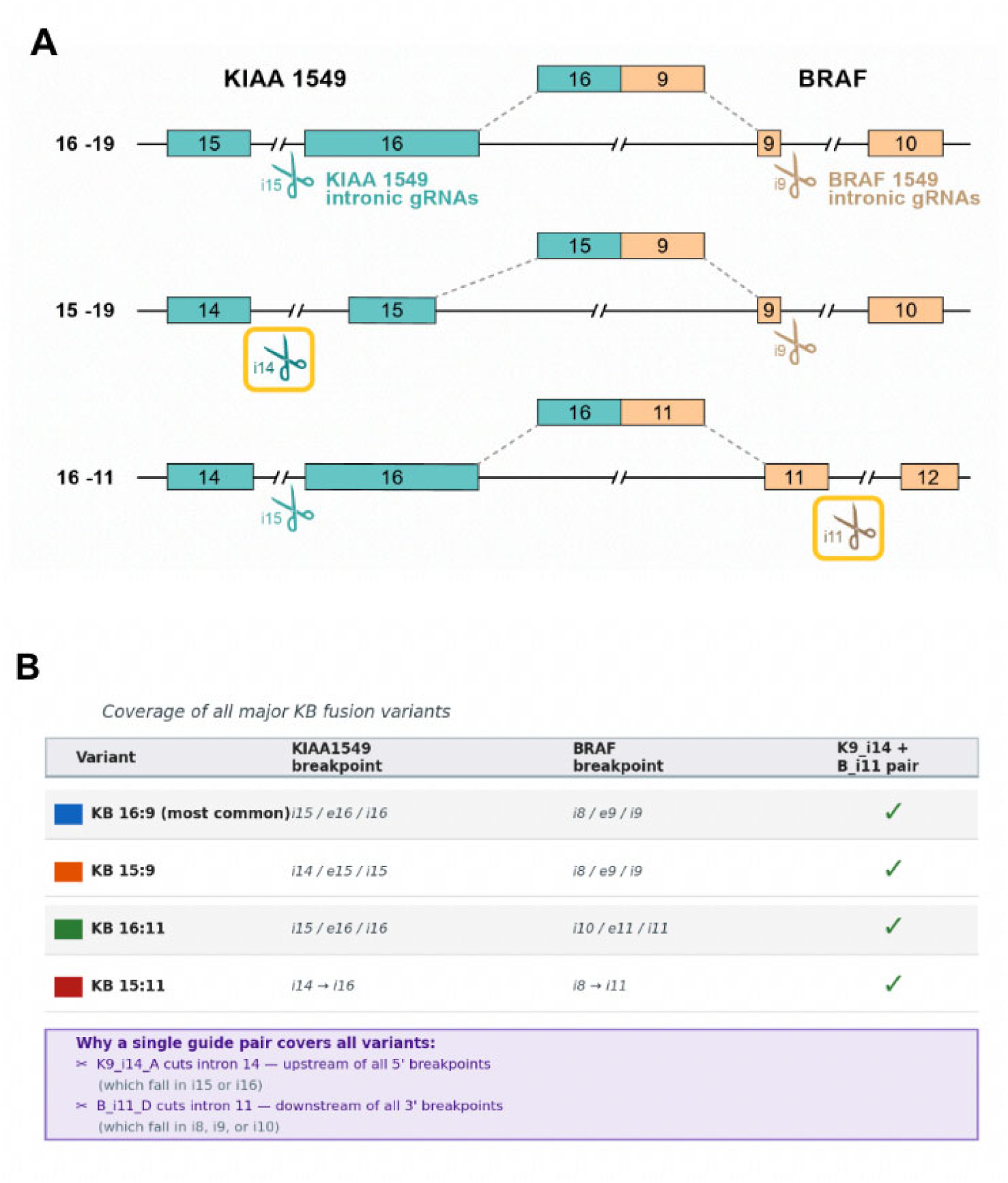
Double-cut intronic excision strategy for KIAA1549—BRAF. (A) Schematic of the 7q34 tandem duplication showing positions three major KB fusion variant breakpoints relative to KIAA1549 introns 14/15 and BRAF introns 9/11. Guide RNA pair targeting KIAA1549 intron 14 (left guide, yellow box) and BRAF intron 11 (right guide, yellow box); dual-cut excision removes the fusion junction and is expected to silence the oncogene. (B) Coverage of all four major fusion variants by the K9_i14 + B_i11 guide pair.

**Figure 2.**
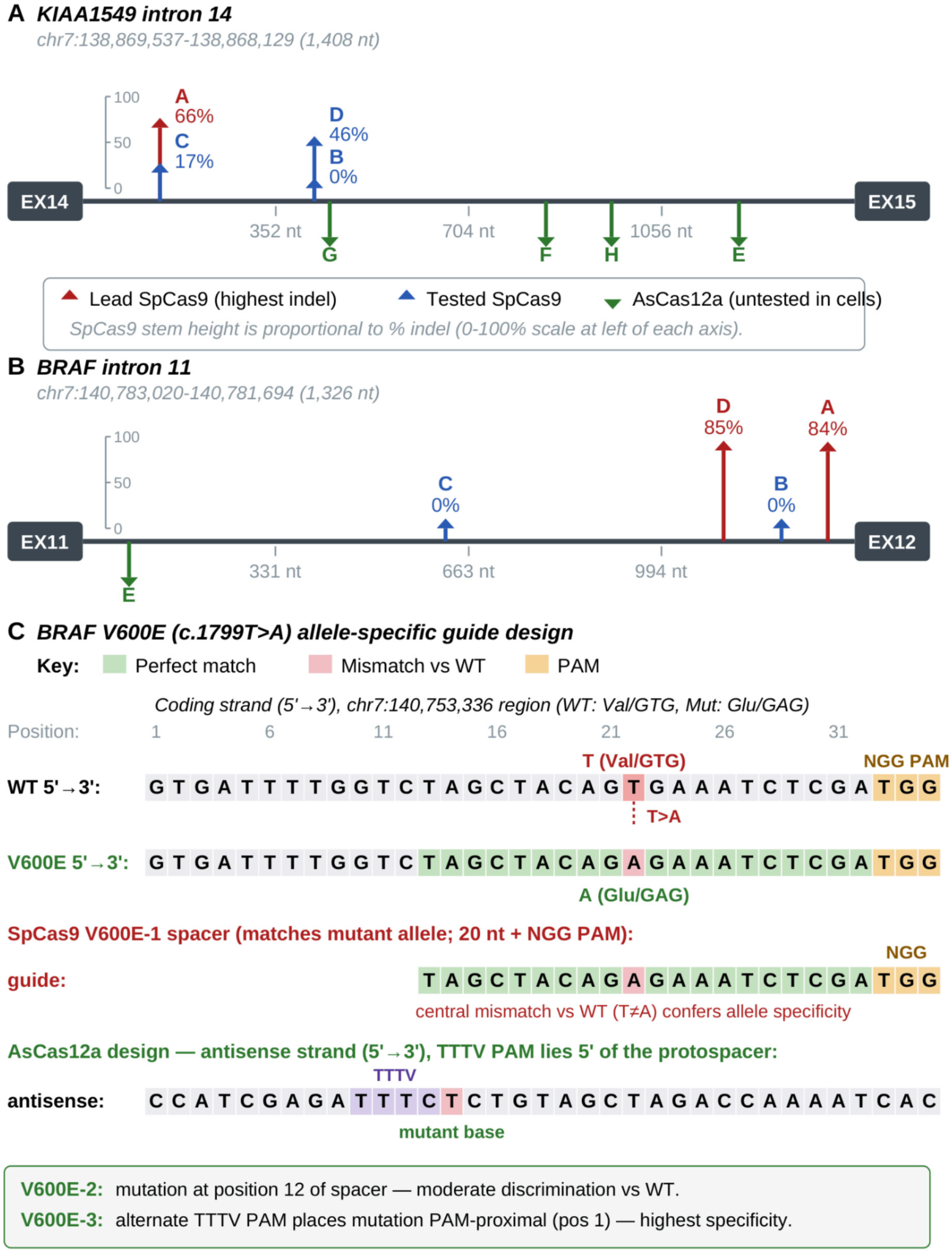
*In silico* design of intronic and allele-specific guide RNAs. (A) Schematic of KIAA1549 intron 14 (chr7:138,869,537—138,868,129) with K9_i14 candidate cut sites annotated. (B) BRAF intron 11 (chr7:140,783,020—140,781, 694) with B_i11 candidates annotated. (C) BRAF V600E locus (c.1799T>A): wild-type vs. mutant sequence, PAM positions, and allele-specific guide design for SpCas9 (NGG PAM on mutant non-template strand) and AsCas12a (TTTV PAM).

**Figure 3.**
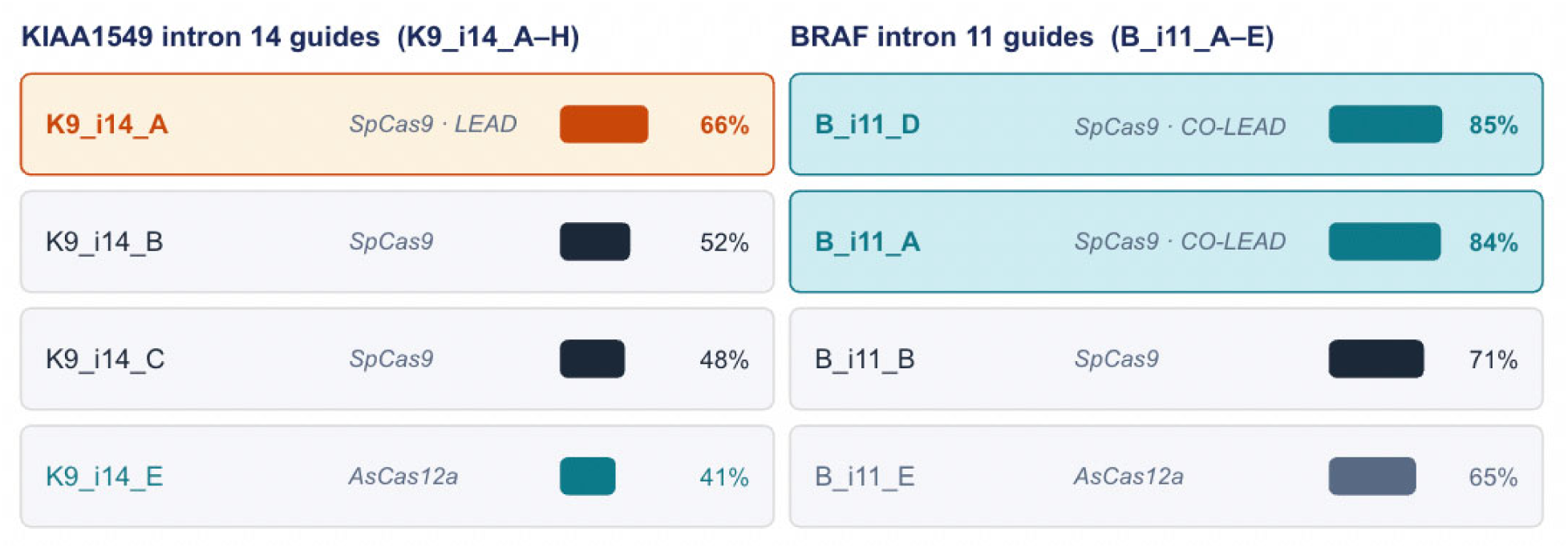
sgRNA activity screen for KIAA1549 intron 14 and BRAF intron 11 guides. ICE indel frequency (%) for each candidate guide in A375 cells. Top performers: K9_i14_A (66%), B_i11 _A (84%), B_i11 _D (85%).

For BRAF V600E, allele-specific SpCas9 guides were screened in A375 cells (homozygous V600E) and 3635 PXA cells (heterozygous V600E). The top V600E allele-specific guide achieved approximately 57% indel frequency in A375 cells, confirming efficient editing at the mutant allele (Fig. 4). In 3635 PXA patient-derived glioma cells, editing efficiency reached approximately 74% by ICE at the V600E locus. Allele specificity was confirmed by sequencing in wild-type BRAF-expressing cells, where editing at codon 600 was not observed at detectable levels, consistent with the absence of a suitable PAM on the wild-type allele.

**Figure 4.**
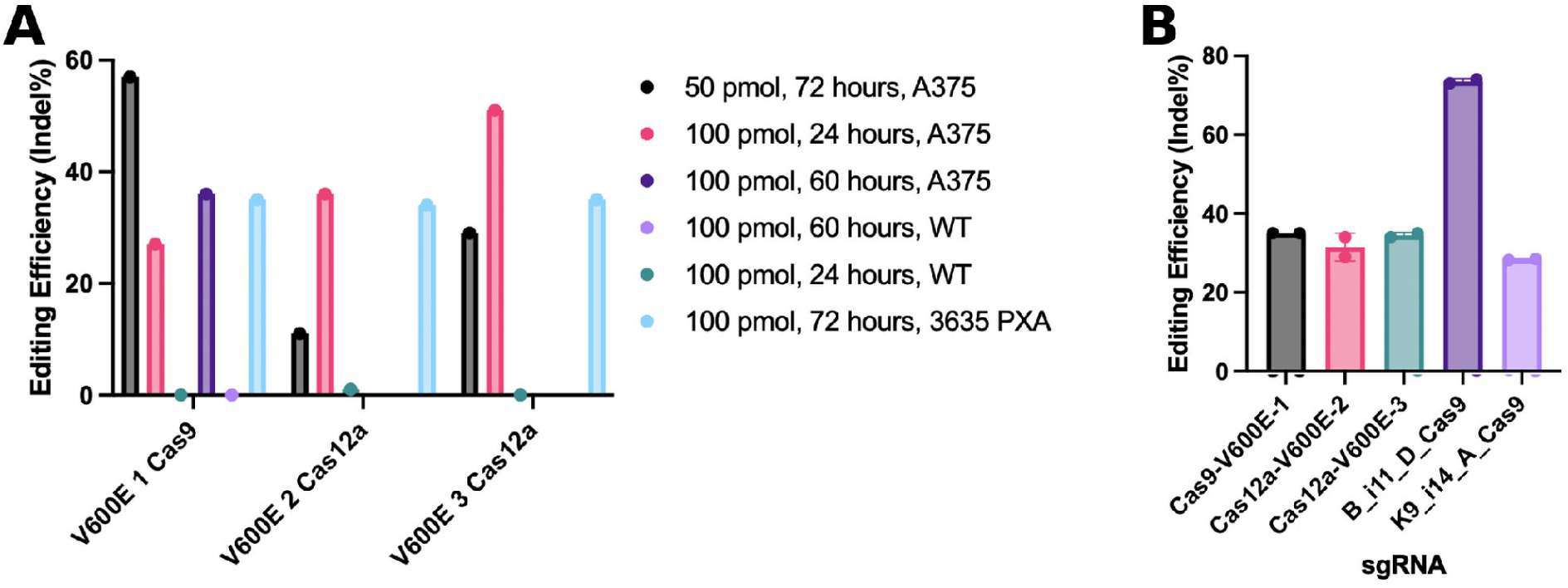
BRAF V600E allele-specific guide RNA screen. (A) ICE indel frequencies for candidate allele-specific SpCas9 guides in A375 cells (V600E homozygous). (B) Editing in 3635 PXA patient-derived glioma cells (V600E heterozygous): ∼74% editing at the V600E locus by the lead guide. (C) Allele specificity confirmation: wild-type BRAF cells show no detectable editing at codon 600

### Dual-cut excision of the KIAA1549—BRAF fusion

Demonstration of productive dual-cut excision requires not only that both guide RNAs cut at high efficiency individually, but also that simultaneous delivery of both RNPs results in the coordinated double-strand breaks necessary for excision rather than sequential single-cut indel formation. To test this directly, we co-nucleofected K9_i14_A_Cas9 and B_i11 _D_Cas9 RNPs simultaneously into DKFZ-BT-31 7 patient-derived pLGG cells with a molecularly confirmed KB fusion (KB 16:9, KIAA1549 exon 16 — BRAF exon 9). RNPs were delivered to cells at equal molar concentrations and assessed editing outcomes by both sequencing at each individual cut site and by a gain-of-function PCR assay specifically designed to detect the excision product.

The gain-of-function PCR excision assay was designed with a forward primer anchored within KIAA1549 intron 14 downstream of the K9_i14_A cut site, and a reverse primer within BRAF intron 11 upstream of the B_i11 _D cut site. In the unedited genome — where the two intronic regions are separated by approximately 1.9 megabases of duplicated chromosomal sequence — no amplicon is produced because the primers are too far apart for conventional PCR amplification. Following successful dual-cut excision and NHEJ-mediated joining of the two intronic ends, the primer pair flanks a predicted amplicon of approximately 1000 base pairs spanning the novel KIAA1549 intron 14 / BRAF intron 11 junction. This amplicon is therefore diagnostic of productive excision and is absent in cells where only single-cut indels have occurred at either site.

### Evidence for a KIAAl549—BRAF fusion in patient-derived WUPA pilocytic astrocytoma cells and a primer-diving strategy for breakpoint localization

To evaluate the KIAA1549—BRAF dual-cut editing strategy in a disease-relevant cellular model, we obtained WUPA8 and WUPA11, patient-derived pilocytic astrocytoma cells from the Gutmann Laboratory at Washington University in St. Louis. These two lines originate from pediatric PA tumors. As a comparator with a molecularly confirmed KB fusion, we included DKFZ-BT-31 7 cells (KB 16:9, KIAA1549 exon 16 — BRAF exon 9).

#### sgRNA activity in WUPAll cells

Prior to dual-cut excision experiments, individual guide RNA activity at the KIAA1549 intron 14 and BRAF intron 11 target sites was assessed by ICE analysis following RNP nucleofection in WUPA11 cells. B_i11 _D_Cas9 achieved 88% indel frequency (KO score 81) and B_i11 _A_Cas9 achieved 86% (KO score 80), consistent with A375 results. K9_i14_A_Cas9 achieved 41 % indel frequency (KO score 41) in WUPA11 cells, somewhat lower than the 66% in A375, likely reflecting differences in chromatin accessibility or cell cycle distribution. These data confirm that both target sites are accessible to Cas9 in patient-derived pLGG cells (Fig. 6).

Dual-cut editing produces excision and inversion products in WUPA cells. To test whether the K9_i14_A and B_i11 _D guide pair could generate the expected editing outcomes, we delivered both RNPs by nucleofection and assessed outcomes by gain-of-function PCR. In cells that received both guides, we detected excision products — NHEJ-mediated rejoining of the K9_i14 and B_i11 cut ends — and inversion products at both the 5’ and 3’ boundaries of the rearranged segment. Both were also detected in DKFZ-BT-31 7 cells (confirmed KB 16:9). No bands were detected in untreated negative controls (Fig. 5).

**Figure 5.**
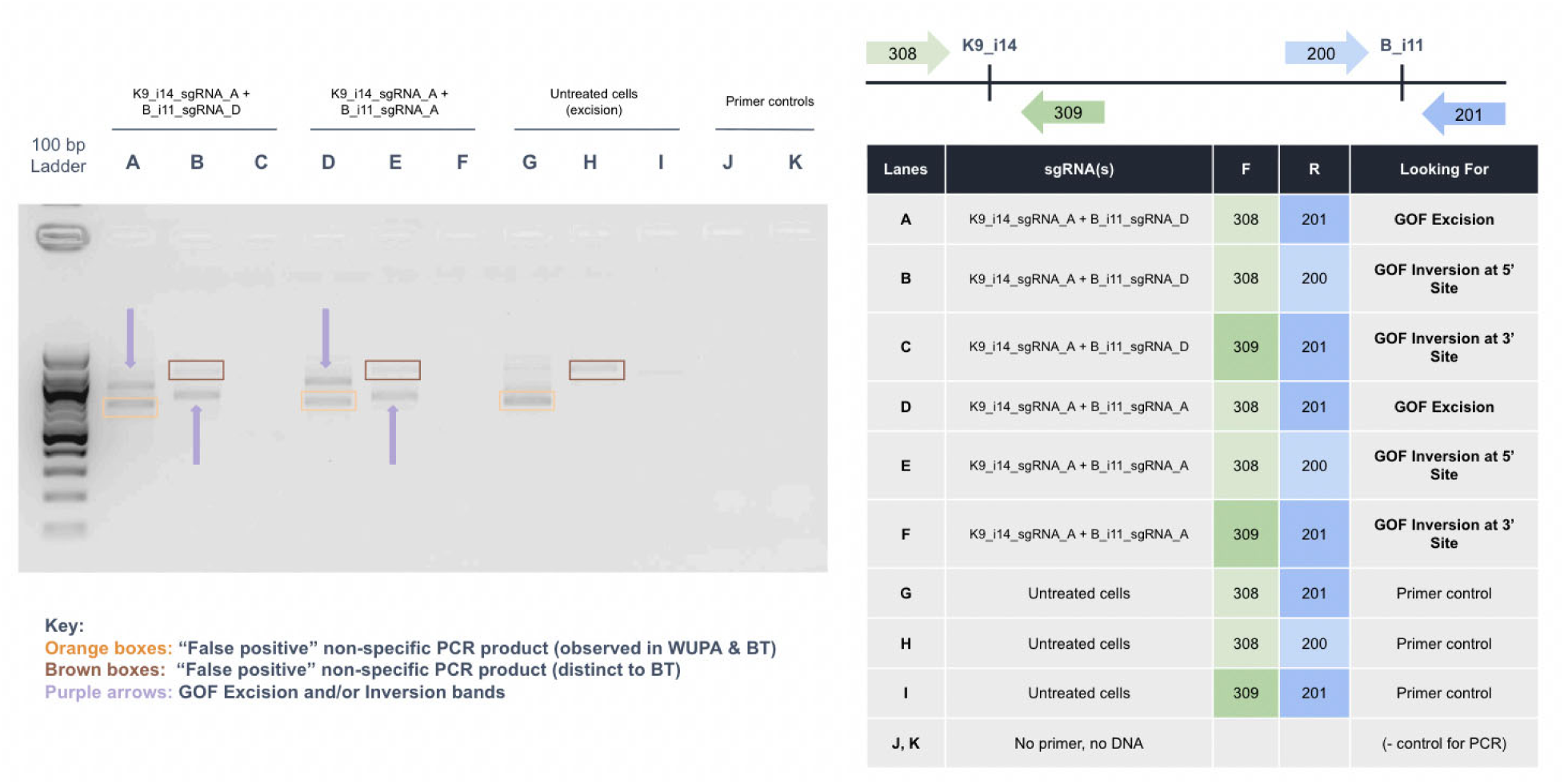
Dual-cut excision of the KIAAl549—BRAF fusion junction assessed by gain-of-function PCR. Gel electrophoresis of GOF PCR amplicons from cells co-nucleofected with K9_i14_A + B_i11 _D (dual guide) vs. single-guide controls and untreated cells. The excision junction band (purple arrow) is present only in the dual-guide condition.

**Figure 6.**
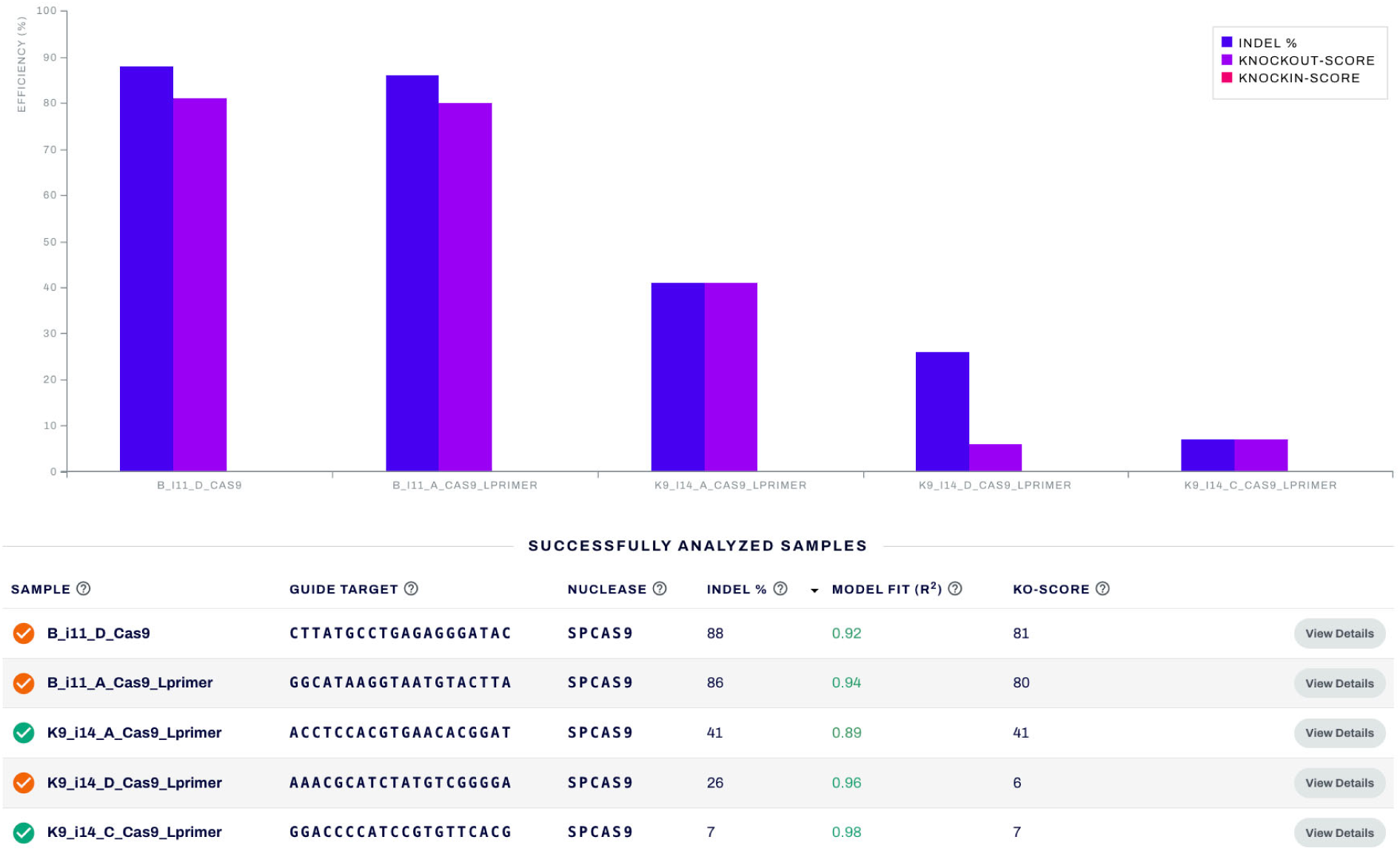
sgRNA activity in WUPAII patient-derived pilocytic astrocytoma cells. ICE indel frequency and KO score for B_i11 _D_Cas9 (88%, KO 81), B_i11 _A_Cas9 (86%, KO 80), K9_i14_A_Cas9 (41 %, KO 41), K9_i14_D_Cas9 (26%), and K9_i14_C_Cas9 (7%) following RNP nucleofection. Data from Synthego ICE analysis.

**Figure 7.**
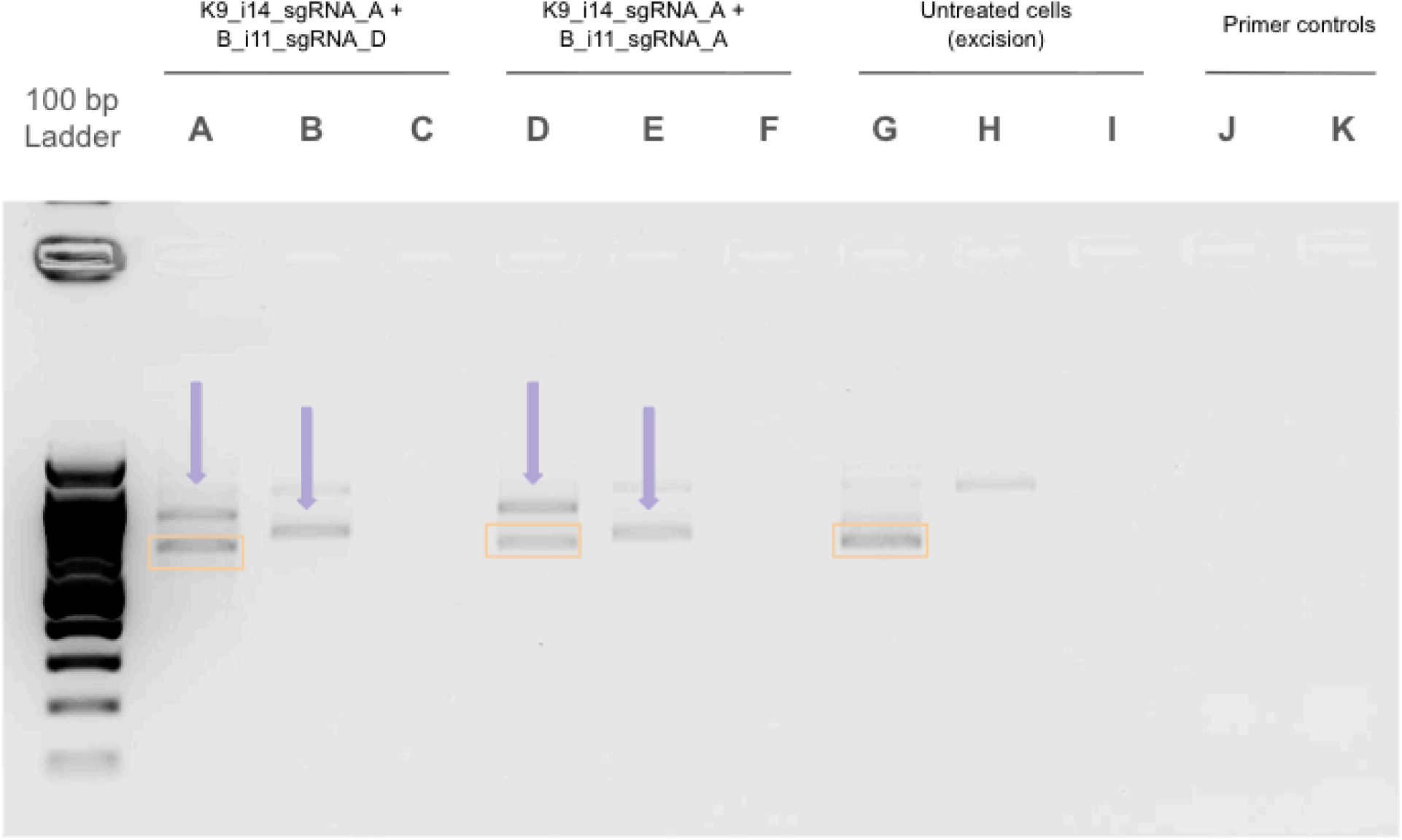
Gain-of-function PCR in WUPAII cells. (A) WUPA11 cells nucleofected with K9_i14_A + B_i11 _D RNPs (dual guide), single-guide controls, and untreated cells. Purple arrows: excision and inversion bands present only in dual-guide conditions. Orange boxes: non-specific alternative amplification. DKFZ-BT-31 7 cells (confirmed KB 16:9 fusion) show an equivalent band pattern.

The detection of inversion amplicons is particularly significant. As described in Materials and Methods, below, 5’ and 3’ inversion bands can only be generated if the KIAA1549—BRAF tandem duplication is present: these amplicons span the junction between the inverted duplicated segment and its flanking chromosomal context, a junction absent in an unfused genome. The GOF PCR results — excision band, 5’ inversion band, and 3’ inversion band, all absent in untreated cells and recapitulated in DKFZ-BT-31 7 — provide strong PCR-level evidence that WUPA cells harbor a KB fusion-containing allele susceptible to dual-cut editing.

Long-read sequencing does not resolve the fusion breakpoint. Oxford Nanopore whole-genome long-read sequencing of WUPA genomic DNA did not identify a canonical KIAA1549—BRAF junction-spanning read at any of the breakpoint intervals of the four major KB fusion variants. Possible explanations include an atypical breakpoint outside the canonical variant intervals, a breakpoint within a highly repetitive intronic region resistant to long-read mapping, or a low variant allele fraction reducing junction-spanning read capture. We regard the GOF PCR evidence — excision and inversion bands reproduced across two independent WUPA lines and mirrored in DKFZ-BT-31 7 — as evidence that a KB fusion is present, and interpret the ONT result as a potential failure to detect rather than conclusive evidence of its absence.

## DISCUSSION

This study describes the design and experimental validation of a CRISPR guide RNA strategy for the two primary oncogenic drivers of pediatric low-grade glioma: the KIAA1549—BRAF oncofusion and the BRAF V600E point mutation. The central design achievement is a ‘catch-all’ intronic excision guide pair — K9_i14_A targeting KIAA1549 intron 14 and B_i11 _D targeting BRAF intron 11 — that addresses all four major KB fusion variants with a single set of reagents.

This approach circumvents the major translational barrier of pLGG heterogeneity: across the hundreds of children diagnosed with KB fusion-positive pLGG each year, the specific variant is detected by molecular sequencing and currently informs clinical classification, but would also complicate a variant-specific therapeutic strategy. By targeting intronic sequences that are invariantly present in all four variants, we eliminate the need for patient-specific guide design while maintaining the precision required for allele-specific editing.

B_i11 _A and B_i11 _D reached 84—85% indel frequency in A375 cells, and the V600E lead guide achieved approximately 57% in A375 and 74% in patient-derived 3635 PXA glioma cells. These values, obtained by RNP nucleofection (which provides an upper-bound estimate of guide RNA activity in the absence of delivery-related inefficiencies), compare favorably to the less than 20% productive outcomes reported in the EDIT-1 01 trial and demonstrate that the target sites identified in this study are genuinely accessible to Cas9. The difference between V600E editing in A375 (57%) and 3635 PXA (74%) likely reflects differences in chromatin accessibility and cell cycle distribution between the two lines rather than guide RNA-intrinsic factors; the higher editing in patient-derived glioma cells is clinically relevant and encouraging.

The gain-of-function PCR excision assay represents a critical methodological contribution of this study. Unlike ICE analysis, which detects indels at individual cut sites without distinguishing between single-cut and dual-cut editing outcomes, the junction-spanning gain-of-function PCR amplicon is produced only by genuine dual-cut excision or inversion followed by NHEJ rejoining of the correct ends. The 191 bp amplicon design places the forward primer within KIAA1549 intron 14 (downstream of the K9_i14_A cut) and the reverse primer within BRAF intron 11 (upstream of the B_i11 _D cut); in the unedited genome, these primer sites are separated by approximately 1.9 Mb and produce no amplicon. This assay provides a direct, qualitative readout of the therapeutically relevant editing outcome, and can be deployed in any cell type or *in vivo* tissue sample that contains the KB fusion duplication.

The allele-specificity of the V600E editing strategy deserves particular emphasis in the context of therapeutic safety. Wild-type BRAF is an essential signaling kinase in neurons, astrocytes, and other CNS cell types, where it mediates activity-dependent transcriptional programs, survival signaling, and synaptic plasticity. Non-allele-specific disruption of BRAF in CNS cells would be expected to cause severe neurological dysfunction. The PAM-based allele specificity of the V600E guide strategy — targeting a PAM present exclusively on the mutant allele — provides a molecular safeguard: wild-type BRAF-expressing cells simply lack the recognition element for the guide RNA. This is qualitatively different from the non-specificity of pharmacological RAF inhibitors, which inhibit both wild-type and V600E BRAF kinase activity and must rely on the differential sensitivity of oncogene-addicted versus normal cells to achieve a therapeutic window.

Several important caveats limit the conclusions that can be drawn from the present experiments. The A375 melanoma cell line, while a standard substrate for CRISPR guide RNA screening and V600E editing validation, is not a CNS cell and does not carry the KB fusion; it was used primarily as a tractable model for assessing guide RNA activity at the target loci. The 3635 PXA line provides more disease-relevant context but is a V600E-driven tumor, not a KB fusion tumor; dedicated KB fusion-positive pLGG cell lines are scarce, and validation of the K9_i14 + B_i11 dual-cut strategy in a genuine KB fusion-expressing pLGG cell line remains an important pending experiment. We look to pursue this validation in pilocytic astrocytoma lines, which harbor the KB 16:9 fusion and grow in immunodeficient murine hosts, enabling both *in vitro* and *in vivo* testing.

The most critical next step for the KB fusion editing strategy is confirmation that productive excision - as measured by the gain-of-function PCR junction assay - is achievable at therapeutically relevant frequencies in pLGG-specific cell types. Validation of the functional consequences of excision remains an important open question. Proliferation, viability, and colony formation assays comparing dual-guide edited vs. unedited cells would directly test whether removal of the KIAA1549-BRAF oncofusion is sufficient to suppress the growth advantage it confers. Western blot quantification of fusion protein reduction using an anti-BRAF antibody targeting the C-terminal kinase domain would provide direct molecular evidence of oncogene disruption; however, this assay was not completed in the current work because patient-derived pLGG cells grow too slowly to yield sufficient protein lysate for reliable immunoblotting, a known technical limitation of primary glioma cultures. Phospho-MEK and phospho-ERK as downstream readouts of BRAF kinase activity represent additional future experiments that would directly link excision efficiency to pathway suppression. Looking beyond cell culture validation, *in vivo* delivery of the KIAA1549-BRAF excision guides represents the natural translational endpoint for this work and would allow us to assess its efficacy in a disease model.

## MATERIALS AND METHODS

### Cell lines and culture

A375 human melanoma cells (ATCC CRL-1619; BRAF V600E homozygous) were maintained in Dulbecco’s Modified Eagle’s Medium (DMEM) supplemented with 10% fetal bovine serum (FBS) and 1 % penicillin/streptomycin (pen-strep) at 37°C, 5% CO_2_. Cells were passaged at 70-80% confluency using 0.25% trypsin-EDTA. 3635 PXA patient-derived pleomorphic xanthoastrocytoma cells (BRAF V600E heterozygous) were maintained in Dulbecco’s Modified Eagle’s Medium (DMEM) supplemented with 10% fetal bovine serum (FBS) and 1 % penicillin/streptomycin (pen-strep) at 37°C, 5% CO_2_. Cells were passaged at 70-80% confluency using 0.25% trypsin-EDTA. All cell lines were tested for mycoplasma contamination using the MycoAlert PLUS detection kit (Lonza) prior to experiments. A summary of cell sources is provided in Table 2.

**Table 2.** Cell line summary: genotype, source, culture conditions, and nucleofection parameters.

|  |  |  |
| --- | --- | --- |
| **DKFZ-BT Cells** | Pilocytic Astrocytoma Cells<br>*Till Milde Lab* | Has KB fusion; immortalized with SV40 |
| **WUPA11 Cells** | Pilocytic Astrocytoma Cells<br>*Gutmann Lab* | Presumed KB fusion cell line based on gain-of-function PCR and gel electrophoresis results; long-read sequencing attempt did not detect common KB fusion |
| **A375 Cells** | Human Melanoma Cells<br>*UC Berkeley Cell Culture Facility* | Harbors V600E mutation; non-KB fusion cell line |

### *In silico* guide RNA design and analysis

Candidate sgRNA spacers for KIAA1549 intron 14 (hg38: chr7:138,869,537-138,868,129, reverse strand) and BRAF intron 11 (hg38: chr7:140,783,020-140,781, 694) were identified by submitting each intron sequence to CHOPCHOP v3 (https://chopchop.cbu.uib.no/)^22^ using the human genome (hg38) as the reference for intronic coordinates, SpCas9 (NGG PAM) as the nuclease, and selecting candidates within the defined target window. Predicted on-target activity (Doench 2016 score) and MIT specificity score were computed for all candidates. For BRAF V600E allele-specific design, a 100-nucleotide window centered on codon 600 (c.1799) was submitted to CHOPCHOP in both wild-type and V600E sequence contexts; guides uniquely supported by a PAM in the V600E context but not in the wild-type context were prioritized. All candidates were evaluated for off-target prediction in the hg38 genome up to four mismatches; candidates with predicted exonic off-targets with two or fewer mismatches were excluded.

### RNP preparation and nucleofection

sgRNA oligonucleotides were synthesized by IDT (Alt-R sgRNA format; 2 nmol scale) and resuspended at 100 pM in IDT Nuclease-Free Duplex Buffer. RNP complexes were assembled by mixing annealed sgRNA with Aldeveron 0C-Cas9-triNLS (10 pM) at a 1.2:1 molar ratio in RNP formation buffer (NaK, NaCl, trehalose, and 0.1 % MgCl_2_) and incubating at room temperature for 10 minutes.

For single-guide screens, one RNP per well was nucleofected. For dual-guide excision experiments, two RNPs (K9_i14_A_Cas9 + B_i11 _D_Cas9) were assembled independently, mixed at equimolar ratio, and co-nucleofected. A375 cells (200,000 cells per reaction) or 3635 PXA cells (200,000 cells per reaction) were nucleofected using the Lonza SF Kit 4D-Nucleofector per the manufacturer’s protocol. Cells were recovered in pre-warmed DMEM + 10% FBS and plated in 48-well format.

### ICE analysis

Genomic DNA was extracted 48—72 hours post-nucleofection using QuickExtract DNA Extraction Solution (Lucigen). PCR amplicons (150—200 bp on each side of each cut site) were generated using the primer pairs described in Table 3. Sanger sequencing was performed at the UC Berkeley DNA Sequencing Facility. Indel frequency and KO score were calculated using Synthego ICE v3 (https://ice.synthego.com).

**Table 3.**
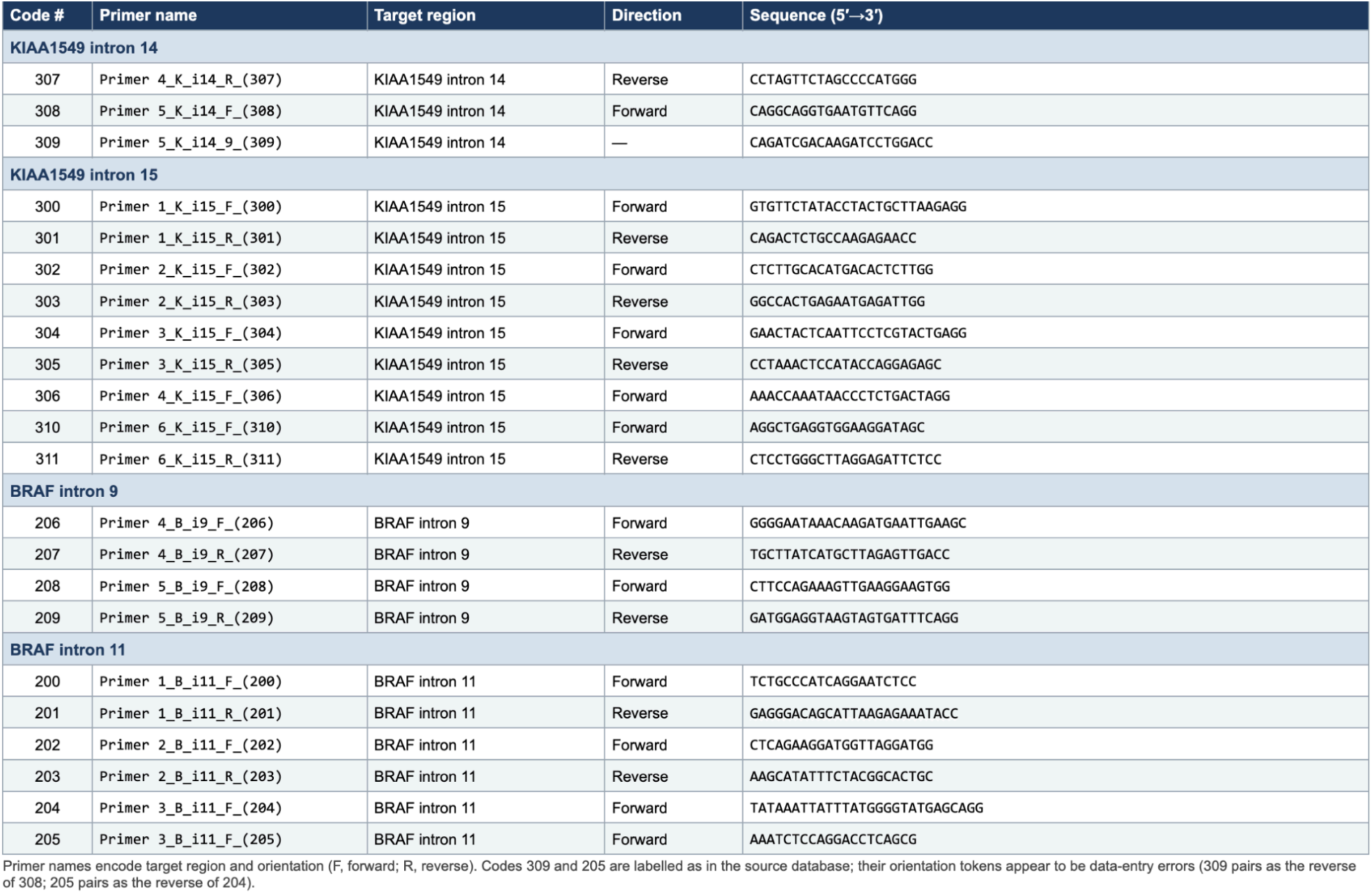
PCR primer sequences (GOF PCR primers: 308, 200, 309, 201); PCR cycling conditions (PrimeStar GXL, 67°C, 1% agarose); expected amplicon sizes for excision and inversion products.

### Gain-of-function PCR excision and inversion assay

A gain-of-function (GOF) PCR assay was developed to detect productive dual-cut editing outcomes — specifically excision and inversion of the KIAA1549—BRAF fusion segment — in a manner that is completely silent in unedited cells regardless of fusion status. No amplicon can be produced from any primer combination in the absence of Cas9 editing: the primers flanking the K9_i14_A and B_i11_D cut sites are separated by approximately 30 kilobases in the fused genome. A detectable band arises only when editing rearranges the two cut ends into close physical proximity, either by excision (direct rejoining of the KIAA1549 and BRAF intronic ends) or by inversion (re-insertion of the excised segment in reversed orientation).

Four primers were positioned across the two cut sites. Two flank the K9_i14_A cut in KIAA1549 intron 14: primer 309 (upstream) and primer 308 (downstream). Two flank the B_i11_D cut in BRAF intron 11: primer 200 (upstream) and primer 201 (downstream). Three amplicon classes become detectable after dual-cut editing: (1) Excision band — primers 308 + 201 face each other across the excision junction, produced by NHEJ rejoining (c191 bp). (2) 5’ inversion band — when the segment is inverted, the 5’ junction brings primers 309 + 200 into proximity. (3) 3’ inversion band — the 3’ junction is detected by a distinct 308 + 201 amplicon. Critically, inversion bands are only possible if the KIAA1549—BRAF tandem duplication is present; in an unfused genome the cut sites are separated by 1.9 Mb and inversion produces no detectable amplicon. Both excision and inversion permanently disrupt the fusion reading frame and are therapeutically relevant outcomes.

Genomic DNA was extracted using QuickExtract (Lucigen) at 48-72 hours post-nucleofection. PCR (50 pL): Takara Bio PrimeStar GXL with primers 308, 200, 309, 201 at 10 pM each, 200—450 ng genomic DNA, 67°C annealing, 1 min/kb extension. Gel: 2% agarose TAE, 100 bp ladder, Bio-Rad Gel Doc imaging. Outcomes scored by presence of excision and/or inversion amplicons absent in untreated and no-template negative controls.

### WUPA cell sequence identification

WUPA8 and WUPA11 are patient-derived pilocytic astrocytoma cell lines obtained from the Gutmann Laboratory at Washington University in St. Louis. Cells were nucleofected with K9_i14_A_Cas9 and B_i11 _D_Cas9 RNPs as described above, and assessed by ICE and gain-of-function PCR as described above. DKFZ-BT-31 7 cells (confirmed KB 16:9) were included as a positive control.

Oxford Nanopore Technologies (ONT) long-read whole-genome sequencing was performed on WUPA genomic DNA using the ONT Ligation Sequencing Kit on a MinION flow cell. Basecalling and alignment to hg38 used Guppy and Minimap2. Structural variant calling used Sniffles2. The 7q34 locus (chr7:138,000,000-141,500,000) was inspected in IGV for reads spanning KB fusion breakpoint intervals in KIAA1549 introns 14-16 and BRAF introns 9-11.

## ACKNOWLEDGEMENTS

This work was supported by funding from Lindonlight. We are grateful to David Gutmann and members of his laboratory at Washington University in St. Louis, as well as to the DKFZ (German Cancer Research Center), for generously providing the patient-derived pLGG cells used in this study. We thank Scott Geller and the UC Berkeley DNA Sequencing Facility for Sanger sequencing support. We thank the UC Berkeley Cell Culture Facility for their support. We thank Florian Muller and Meng-meng Fu for thoughtful discussions and valuable feedback throughout this work. We thank Trang Duong for assistance with long-read sequencing. We thank Sarah Pyle for assistance with graphical elements of the manuscript.

## WORKS CITED

1. Fangusaro, J. et al. Pediatric low-grade glioma: State-of-the-art and ongoing challenges. Neuro-Oncol. 26, 25–37 (2024).

2. Tian, Y., et al. Detection of KIAA1549-BRAF fusion transcripts in formalin-fixed paraffin-embedded pediatric low-grade gliomas. J. Mol. Diagn. JMD 13, 669–677 (2011).

3. Ostrom, Q. T. et al. CBTRUS Statistical Report: Primary Brain and Other Central Nervous System Tumors Diagnosed in the United States in 2015-2019. Neuro-Oncol. 24, v1–v95 (2022).

4. Fernandez, C. et al. Pilocytic astrocytomas in children: prognostic factors--a retrospective study of 80 cases. Neurosurgery 53, 544–553; discussion 554-555 (2003).

5. Packer, R. J. et al. Carboplatin and vincristine chemotherapy for children with newly diagnosed progressive low-grade gliomas. J. Neurosurg. 86, 747–754 (1997).

6. Merchant, T. E., Conklin, H. M., Wu, S., Lustig, R. H. & Xiong, X. Late effects of conformal radiation therapy for pediatric patients with low-grade glioma: prospective evaluation of cognitive, endocrine, and hearing deficits. J. Clin. Oncol. Off. J. Am. Soc. Clin. Oncol. 27, 3691-3697 (2009).

7. Fangusaro, J. et al. Selumetinib in paediatric patients with BRAF-aberrant or neurofibromatosis type 1-associated recurrent, refractory, or progressive low-grade glioma: a multicentre, phase 2 trial. Lancet Oncol. 20, 1011-1022 (2019).

8. Kilburn, L. B. et al. The type II RAF inhibitor tovorafenib in relapsed/refractory pediatric low-grade glioma: the phase 2 FIREFLY-1 trial. Nat. Med. 30, 207–217 (2024).

9. Bouffet, E. et al. Dabrafenib plus Trametinib in Pediatric Glioma with BRAF V600 Mutations. N. Engl. J. Med. 389, 1108–1120 (2023).

10. Wan, P. T. C. et al. Mechanism of activation of the RAF-ERK signaling pathway by oncogenic mutations of B-RAF. Cell 116, 855–867 (2004).

11. Jones, D. T. W. et al. Tandem duplication producing a novel oncogenic BRAF fusion gene defines the majority of pilocytic astrocytomas. Cancer Res. 68, 8673–8677 (2008).

12. Nobre, L. et al. LGG-16. PREDICTORS OF OUTCOME IN BRAF-V600E PEDIATRIC GLIOMAS TREATED WITH BRAF INHIBITORS: A REPORT FROM THE PLGG TASKFORCE. Neuro-Oncol. 21, ii102–ii102 (2019).

13. Jinek, M. et al. A programmable dual-RNA-guided DNA endonuclease in adaptive bacterial immunity. Science 337, 816–821 (2012).

14. Zetsche, B. et al. Cpf1 is a single RNA-guided endonuclease of a class 2 CRISPR-Cas system. Cell 163, 759–771 (2015).

15. Appay, R., et al. Duplications of KIAA1549 and BRAF screening by Droplet Digital PCR from formalin-fixed paraffin-embedded DNA is an accurate alternative for KIAA1549-BRAF fusion detection in pilocytic astrocytomas. Mod. Pathol. Off. J. U. S. Can. Acad. Pathol. Inc 31, 1490–1501 (2018).

16. Tatevossian, R. G. et al. MAPK pathway activation and the origins of pediatric low-grade astrocytomas. J. Cell. Physiol. 222, 509–514 (2010).

17. Forshew, T. et al. Activation of the ERK/MAPK pathway: a signature genetic defect in posterior fossa pilocytic astrocytomas. J. Pathol. 218, 172–181 (2009).

18. Poulikakos, P. I., Zhang, C., Bollag, G., Shokat, K. M. & Rosen, N. RAF inhibitors transactivate RAF dimers and ERK signalling in cells with wild-type BRAF. Nature 464, 427–430 (2010).

19. Hatzivassiliou, G. et al. RAF inhibitors prime wild-type RAF to activate the MAPK pathway and enhance growth. Nature 464, 431–435 (2010).

20. Haeussler, M. et al. Evaluation of off-target and on-target scoring algorithms and integration into the guide RNA selection tool CRISPOR. Genome Biol. 17, 148 (2016).

21. Lapinaite, A., Doudna, J. A. & Cate, J. H. D. Programmable RNA recognition using a CRISPR-associated Argonaute. Proc. Natl. Acad. Sci. U. S. A. 115, 3368–3373 (2018).

22. Montague, T. G., Cruz, J. M., Gagnon, J. A., Church, G. M. & Valen, E. CHOPCHOP: a CRISPR/Cas9 and TALEN web tool for genome editing. Nucleic Acids Res. 42, W401–W407 (2014).

